# Improving Welfare Through Enrichment: A Case Study in Aged Ex-Laboratory Rhesus Macaques

**DOI:** 10.64898/2026.05.05.719840

**Authors:** Fabrizio Dell’Anna, Valeria Albanese, Roberta Berardi, Michela Kuan, Pier Attilio Accorsi, Giovanna Marliani, Maria Padrell, Miquel Llorente

## Abstract

Rhesus macaques (Macaca mulatta) are widely used as non-human primate models for biomedical research. When housed in captivity, it is essential to provide an environment that supports their natural behaviours; otherwise, they risk developing mood disorders, stereotypies, and other behavioural issues that may lead to physical harm. The objective of this preliminary study was to monitor the behaviour of three aged rhesus macaques (≥ 20 y.o.), relocated from a laboratory to a Rescue Centre for Exotic Animals (Italy), and to assess the impact of novel food enrichments. Behavioural data were collected over 18 weeks, beginning at their arrival, using continuous focal sampling from video recordings. Simultaneously, faecal samples were gathered for cortisol analysis. The study was divided into three phases: a control phase without enrichments, a feeding enrichment phase (divided into two periods), and a final control phase without enrichments. Each phase comprised 900 minutes of observations for each subject. Data were analysed using generalized linear mixed models. Results showed an increase in locomotion during the enrichment and final phase compared to the initial phase. Additionally, a reduction in scratching and body-shaking behaviours was observed in the final phase compared to the initial phase. These findings suggest that implementing an enrichment program can enhance the welfare of aged non-human primates and can be considered a valuable tool in the rehabilitation of non-human primates previously housed in laboratories.

**GRAPHICAL ABSTRACT:** 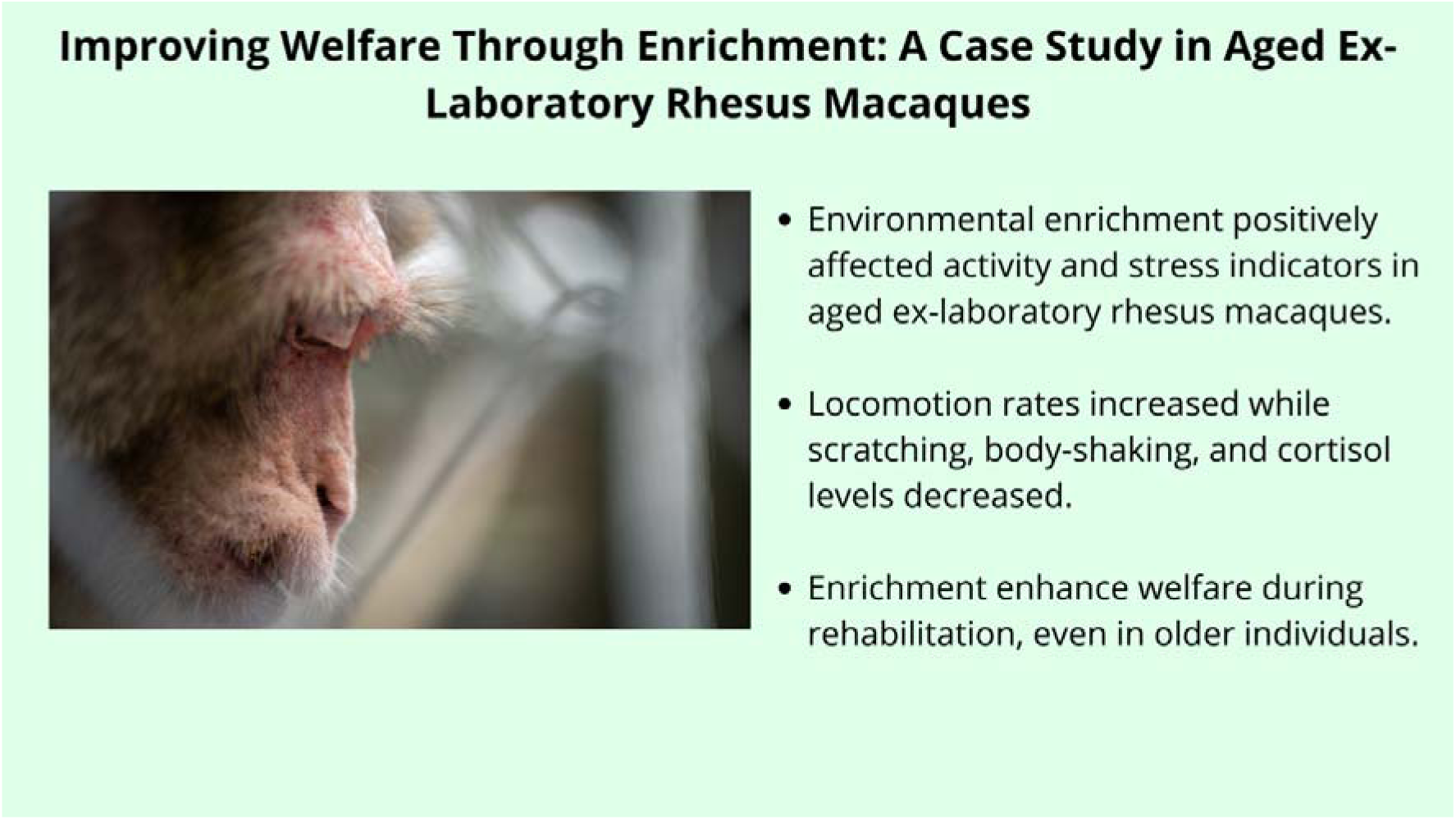

**RESEARCH HIGHLIGHTS:**

- Environmental enrichment positively affected activity and stress indicators in aged ex-laboratory rhesus macaques.
- Locomotion rates increased while scratching, body-shaking, and cortisol levels decreased.
- Enrichment enhance welfare during rehabilitation, even in older individuals.

## INTRODUCTION

Non-human primates (hereafter, primates) are still widely used for experimental purposes in laboratory setting. Globally, over 100,000 individuals are estimated to be used annually in research (Padrell et al., 2021), with *Macaca mulatta* being one of the most common species (Carlsson et al., 2004). The European Commission has issued a highly restrictive report on the topic (Adams et al., 2019), and the Netherlands National Committee for the protection of animals used for scientific purposes (NCad) has recommended the immediate cessation of primate use, citing unsustainability due to ethical, scientific, and legal reasons (Koëter et al., 2016). Nonetheless, primates are still widely used for experimental purposes in Italy, despite strong constraints on their use by national and international regulations and significant ethical opposition to their employment globally (Carvalho et al., 2019; EU Commission Summary Report, 2020; Padrell et al., 2021). Although the Italian Decree no. 73/28-03-2022 provides for the release and rehoming of animals used for experimental purposes, there is no official report on the number of animals rehomed at the end of procedures. Rehoming primate species previously housed under laboratory conditions necessitates a specific environment and a rehabilitation programme to restore their physical and psychological welfare state (Albanese et al., 2021). Laboratory housing conditions, which animals may endure for many years, can cause physical and behavioural changes such as alopecia, and the development of abnormal behaviours, like pacing or hair plucking (Lutz et al., 2022). Behavioural problems commonly observed in captive primates include stereotypic behaviours, self-directed behaviours (e.g., self-mutilation, hair pulling), and aggression towards conspecifics or caretakers. Stereotypic behaviours, such as pacing, rocking, or repetitive movements, are often seen as a coping mechanism in response to the barren and unstimulating abnormal captive environment (Lutz and Novak, 2005). Additionally, boredom and inactivity (i.e., the behavioural state resulting from inadequate environmental stimulation, Lutz and Novak, 2005) can promote the onset of the aforementioned behaviours, which can hinder the rehabilitation of the animals (Cheyne, 2006). However, environmental enrichment techniques effectively provide opportunities for developing natural species-specific behavioural repertoires and ethological needs, reducing the rate of abnormal behaviours, and improving the welfare of animals under human care (Lutz and Novak, 2005; Gronqvist et al., 2013). To ensure effectiveness, enrichment programmes should consider the species-specific natural history, and the behavioural ecology of the species (Lutz and Novak, 2005), but also the individual life experiences and personality profiles (Goswami et al., 2020; Rey et al., 2024). For example, in primates, such as macaques, social housing serves as effective enrichment, and in stressful situations, companions act as a social buffer (Hennessy et al., 2016; Hannibal et al., 2017). Indeed, in their natural environment rhesus macaques (*Macaca mulatta*) live in multimale-multifemale social systems characterised by strong bonds between female kin (Lewis and Prongay, 2015). Providing cognitive challenges and environmental enrichment have shown to be effective solutions to address the behavioural welfare issues faced by captive primates (Clarck and Smith, 2013; Padrell et al. 2022). Examples of effective environmental enrichment include structural elements (such as platforms, ropes, and water pools), nutritional additions (e.g., new food items or puzzle feeders), and manipulative items (toys and novel objects, Cannon et al., 2016). Previous studies on rhesus macaques housed under laboratory conditions, have shown that access to complex outdoor enclosures leads to higher scores of locomotion and a decrease in self-directed oral behaviours (O’Neill et al., 1991). These enrichment types prove useful when it is not possible to keep animals in groups and are effective in reducing the rate of abnormal behaviour (Cannon et al., 2016; Wooddell et al., 2019). Additionally, the evaluation of stressful situations requires consideration of both behavioural parameters and physiological indices (Rushen et al., 2011), such as cortisol concentration. Stressful conditions activate the hypothalamic–pituitary–adrenal (HPA) axis, leading to cortisol release (Moberg, 2000). Consequently, measuring cortisol and its metabolites in faeces is widely employed non-invasive method in animal welfare research to assess stress levels (Palme, 2005, 2012; Schwarzenberger, 2007). Rescue centres, supported by primatologists and veterinarians, accept animals from laboratories and focus on their physical and behavioural rehabilitation (Carlsson et al., 2004). The aim of this study was to evaluate the effect of different types of environmental enrichment on the behaviour and welfare of aged ex-laboratory macaques, to improve rehabilitation programmes for rescued non-human primates. To achieve a multidisciplinary approach, we combined behavioural observations with physiological measures. We developed the following hypotheses and tested the following predictions:

### Hypothesis 1

Enrichments can improve the welfare of aged macaques recovered from laboratories by keeping them engaged, preventing an excess of inactivity.

#### Prediction 1

Individuals spend more time being active (e.g., moving) in the post-treatment phase compared to the pre-treatment phase (prediction 1a). Moreover, individuals will spend more time manipulating objects in the post-treatment phase compared to the pre-treatment phase (prediction 1b).

### Hypothesis 2

Environmental enrichment activities can improve the welfare of aged macaques recovered from laboratories by decreasing the rate of abnormal and anxiety-like behaviours.

#### Prediction 2

The rate of observed abnormal and anxiety-like behaviours (i.e., scratching, body shaking) will be lower in the post-treatment phase compared to the pre-treatment phase.

### Hypothesis 3

Environmental enrichment activities can improve the welfare of aged macaques recovered from laboratories by reducing their physiological stress.

#### Prediction 3

The faecal cortisol concentration will be lower in the post-treatment phase compared to the pre-treatment phase.

## 2. MATERIALS AND METHODS

### Subjects and Study Site

The study subjects comprised three rhesus macaques (*Macaca mulatta*) relocated from the laboratory of the Faculty of Medicine and Surgery at the University of Verona (Italy) to the Animanatura Wild Sanctuary in Semproniano in June 2021. All subjects were males, with two individuals of aged 20 years (born in 2001) and the oldest aged 28 (born in 1993). Detailed information about the study subjects is available in Supplementary Materials S-1.

Each subject’s enclosure was comprised of (i) an outside compartment (29 m^2^) furnished with wooden structures and hammocks made of firehose, including various environmental objects that could have been manipulated (e.g., sticks, stones, etc.), and (ii) an inside compartment (10.5 m^2^) furnished with wooden platforms (see Supplementary Materials S-2). The floor of the indoor compartments was covered with wood chips. In wintertime, an air heating system maintained a stable temperature (15–20 C°) in the inside compartments. The animals were provisioned three times a day with commercial pellets in the morning (Kasper Fauna Food-Primate PT1), fruit and vegetables at midday, and carrots in the afternoon. In addition, nuts and dried fruit were provided at least twice a week.

### Ethical Statement

This project followed the protocols approved by the European Parliament and Council’s Directive 2010/63/EU of 22 September 2010 on the protection of animals used for scientific purposes. It also followed the institutional guidelines for the care and management of primates established by the International Primatological Society and LAV.

### Behavioural Observations

We collected between June and October 2021 and consisted of 352 focal samples collected by VA and RB, by means of 30-minute and 15-minute focal animal samples using continuous sampling (Altmann, 1974) of the three subjects. Thirty-minute focal samples were used in the pre- and post-treatment phases, whereas 15-minute focal samples were used in the treatment phase (15 minutes of continuous observation at the beginning of the distribution of the enrichment, and 15 minutes of observation 45 minutes after the beginning of the distribution of the enrichment). We conducted the observations either during the morning (e.g., before 12pm) or during the afternoon (after 12:00pm), balancing the number of morning and afternoon sessions during the study. During the enrichment phase, the same data collection protocol was used during the days with and without enrichment (see Table 2). Total observation time amounted to 7860 minutes (131 hours) of observation. We video-recorded all experimental sessions, and behaviours later coded from the videos using BORIS software, an application that facilitates the decoding and analysis of animal behaviour (Friard and Gamba, 2016). We later imported data into Microsoft Excel to perform data wrangling and formatting to prepare for data analysis. During observation, we collected the observed behaviours included in the ethogram showed in Table 1. Additionally, we recorded the presence of disturbances during recording sessions (i.e., tourists, workers). Finally, maximum daily temperature during the observation day was later collected from the database of a nearby regional weather station (Semproniano station, lat. 42.731 N, lon. 11.597 E; sir.toscana.it).

**Table 1.** Ethogram employed in the study, based on Nickelson and Lockard, 1978; Maestripieri et al., 1992, Aureli et al., 1999; Pomerantz et al., 2012; Xu et al., 2012; Sha and Hanya, 2013)

| Behaviour | Definition |
| --- | --- |
| Scratching | Vigorous strokes of the hair with harms or leg. |
| Body shake | Rapid shaking of head and shoulders defined as a self-directed behaviour (SDB) in a context of social anxiety |
| Locomotion | Moving to another location by walking, scampering, scurrying or climbing. |
| Object manipulation | Manipulate or transport artificial object with hands or mouth. |
| Abnormal behaviour | Anomalous behaviour including stereotypic pacing and biting/licking the mesh. |

**Table 2.** Summary of the study design and structure.

|  | **Pre-treatment** | **Treatment 1** | **Treatment 2** | **Post-treatment** |
| --- | --- | --- | --- | --- |
| **Dates** | 08.06.2021-09.07.2021 | 27.07.2021-13.08.2021 | 16.08.2021-03.09.2021 | 07.09.2021-18.10.2021 |
| **Duration** | 6 weeks | 3 weeks | 3 weeks | 6 weeks |
| **Number of sessions** | 30 days | 15 days | 15 days | 30 days |
| **Number of enrichment sessions** | None | 7 | 7 | None |
| **Total amount of faecal samples collected** | 36 | 36 | 36 | 33* |

\*3 faecal samples are missing due to the death of one of the subjects

### Inter□Observer Reliability

One additional coder, blind to the goals of the study, scored 15% of the video recorded sessions (N=38) to assess interobserver reliability. We used two-way random effect intraclass correlation coefficient (ICC) for continuous data (e.g., abnormal behaviours, Koo and Li, 2016; Bateson and Martin, 2021) and Cohen’s k for categorical data (e.g., scratching, Bateson and Martin, 2021). The agreement between the two observers was excellent (ICC=0.9, p < 0.001; Cohen’s k=0.84). Analyses were conducted using the “*irr*” package (Gamer et al., 2021) in R (version 4.3.1, R Core Team 2023).

### Experimental Procedure

The study was conducted following a 3-phases procedure: pre-treatment, treatment and post-treatment phase. Pre-treatment phase had a duration of 6 weeks, starting in June 2021 as soon as the subjects arrived at Animanatura Wild Sanctuary. We observed monkeys for 30 days (Mon-Fri) via 30 minutes continuous video observations. Faecal samples were collected on Tuesdays and Fridays, for a total of 12 samples collected for each subject. Treatment phase had a duration of 6 weeks (i.e., a 3-weeks cycle repeated twice), for a total of 14 (7+7) enrichment sessions. Enrichment sessions were distributed following the scheme: Week 1-Tuesday and Thursday, Week 2 Monday, Wednesday and Friday, Week 3: Tuesday and Thursday. Subjects received a different kind of feeding enrichment in each different session. A list of feeding enrichments used in the study can be found in Supplementary Materials S-2. We collected faecal samples on Monday, Tuesday, Thursday, and Friday, for a total of 24 samples collected for each subject. The treatment phase protocol was used for both the days in which enrichment was offered to the subjects and for the days in which enrichment was not offered. Finally, post-treatment phase had a duration of 6 weeks. We observed monkeys for 30 days (Mon-Fri) via 30 minutes continuous video observations. Faecal samples were collected on Tuesdays and Fridays, for a total of 12 samples collected for each subject. Information about experimental phases is summarized in Table 2.

### Faecal sampling and faecal cortisol metabolite assays

We collected faecal samples from the inside enclosure during the morning and a total of 137 samples were analysed. The samples were collected between June 2021 and October 2021. We collected the samples twice a week (Tuesday and Friday) during the pre-treatment and post-treatment phase and four times a week during the treatment phase (Monday, Tuesday, Thursday and Friday). Each sample, which represented a single subject, was stored in a freezer (−18 °C) until delivery to the laboratory at the Department of Medical Veterinary Sciences of the University of Bologna. We determined the faecal cortisol metabolite levels by radioimmunoassays (RIAs). We modified the extraction methodology from Schatz and Palme (2001). Cortisol metabolite assays in faeces were carried out according to Tamanini et al. (1983). Validation parameters of analyses were as follows: sensitivity 0.19 pg/mg, intra-assay variability 5.9%, inter-assay variability 8.7%. Radioactivity was determined using a liquid scintillation beta counter and a linear standard curve, designed ad hoc by a software programme (Motta and Degli Esposti, 1981). All concentrations were expressed in picograms per milligram of faecal matter (pg/mg).

### Data Analyses

To perform statistical analyses, we utilized seven different generalized linear mixed models (GLMM) using the “*glmmTMB*” package (Bolker, 2019) in R (R Core Team, version 3.5.0). In Models 1-6, we compared macaque’s behaviours across phases (pre□treatment, treatment and post-treatment) by using phase as the main predictor. In particular, these models assessed whether phase predicted the occurrence body shake (Model 1), scratching (Model 2), locomotion (Model 3), object manipulation (Model 4), and abnormal behaviours (Model 5). Models 1 and 2 used behaviour frequency (e.g., scratches bouts count in Model 1) as dependent variable, whereas the duration of the focal observation was used as offset variable. Time of observation (morning/afternoon), maximum daily temperature and presence of disturbances were used control variables, whereas subject identity was used as random effect. Models 1-2 were modelled using a Poisson distribution. Models 3-5 were fitted in similar fashion: given the continuous nature of the behaviours observed, the dependent variable used was the proportion of time spent performing such behaviour during the focal observation (i.e., time spent manipulating an object, divided by the total duration of the observation). Similarly to models 1-2, time of observation (morning/afternoon), maximum daily temperature and presence of disturbances were used as control variables, whereas subject identity was used as random effect. The dependent variable used in Models 3-6 was transformed to rescale data points with value “0” (zero) into extremely low values (e.g., 0.0005), because the *glmmTMB* package cannot handle “0” or “1” values in beta distribution models (Brooks et al., 2017). Models 1-6 were fitted using a dataset that comprised all focal observation (i.e., one data-point is represented by one 15-minutes or 30-minutes focal observation, N=352). Finally, in model 6 we compared faecal cortisol concentration across phases (pre□treatment, treatment and post-treatment) by using phase as the main predictor. Time of observation (morning/afternoon), maximum daily temperature and presence of disturbances were used control variables, whereas subject identity was used as random effect. Model 7 was fitted using a negative binomial distribution. The dataset on which this model is based comprised of 133 datapoints. We excluded cortisol data collected on subject Charlie regarding October 2021 to avoid noise in the data, given to the death of the subject occurred on October 7^th^, 2021. Therefore, three faecal samples pertaining to the subject were excluded from this analysis. We tested whether each model was significantly different from the corresponding null model, which only included control variables and random factors, via likelihood ratio tests (Dobson and Barnett, 2018). In those cases where the predictor had a significant effect, we conducted post-hoc tests using the “*emmeans*” package (Lenth, 2022), to perform Tukey’s multiple pairwise comparisons (p < 0.05), using robust standard errors. In order to test our predictions, we specifically compared behaviours between the pre-treatment phase and the post-treatment phase. Continuous variables were z-transformed to improve model convergence. We used the Variance Inflation Factor (VIF) to examine collinearity across predictor and control variables (Field, 2005). No models were affected by collinearity issues (e.g., VIF< 3), except Model 5 which showed a max VIF value of 3.37. We tested for model overdispersion and, if overdispersion was detected, (e.g., Model 5) we transformed the response variable into a count variable, therefore using focal observation duration as offset variable. Such models were fitted using a negative binomial distribution. All the models presented below were found to not be overdispersed.

## 3. RESULTS

On a descriptive level, results showed an increase in locomotion during the enrichment and final phase compared to the initial phase. On average, subjects moved for only 7% of the time in the initial phase, whereas they were observed moving for 11% of the time in the final phase. Similarly, we found a decrease in resting behaviour in both the enrichment (12% of observation time) and final control phase (13%) compared to the initial phase (24%). In Model 1 (body shake) the full vs. null model comparison was significant (χ2=8.5089, df=3, p=0.037) and post□hoc comparisons indicated that body shake was less likely to occur in the post-treatment phase compared to the pre-treatment phase (β = -0.7541, SE= 0.295, p=0.05). In Model 2 (scratching) the full vs. null model comparison was significant (χ2=32.479, df=3, p<0.001) and post□hoc comparisons indicated that scratching was less likely to occur in the post-treatment phase compared to the pre-treatment phase (β = -0.493, SE= 0.171, p=0.02, Figure 1).

**Figure 1.**
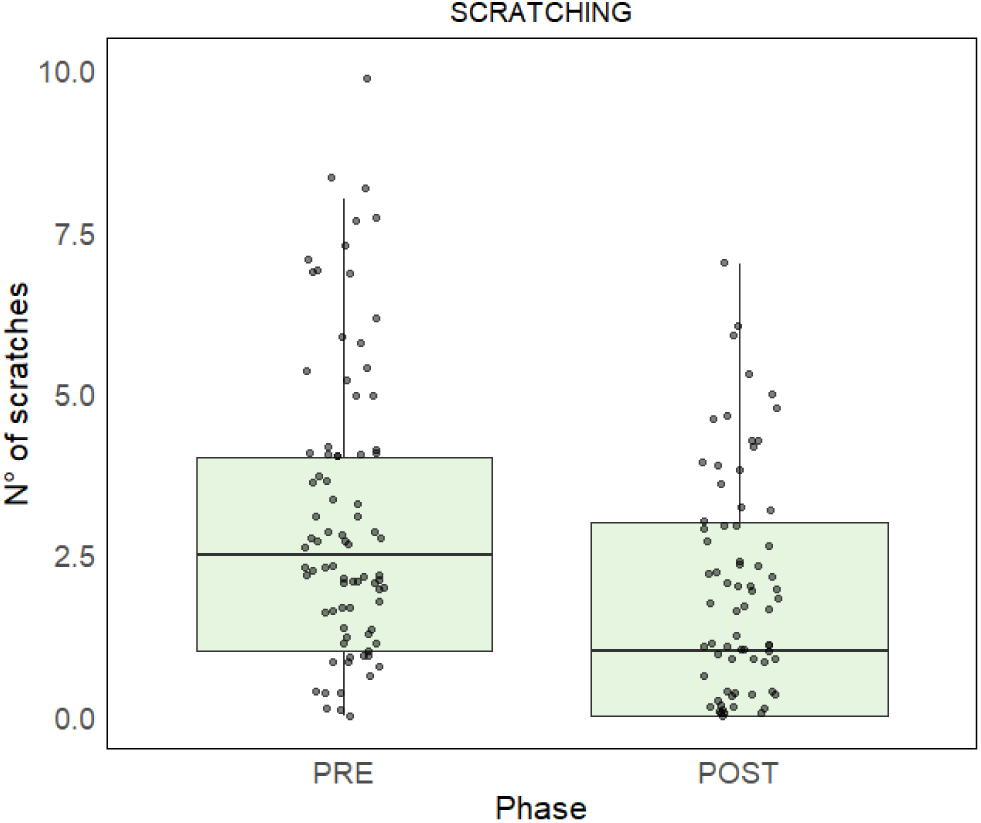
Number of scratches in pre-treatment and post-treatment phases.

In Model 3 (locomotion) the full vs. null model comparison was significant (χ2= 21.55, df=3, p<0.001) and post□hoc comparisons indicated that locomotion was more likely to occur in the post-treatment phase compared to the pre-treatment phase (β = 0.4934, SE= 0.150, p=0.006, Figure 2).

**Figure 2.**
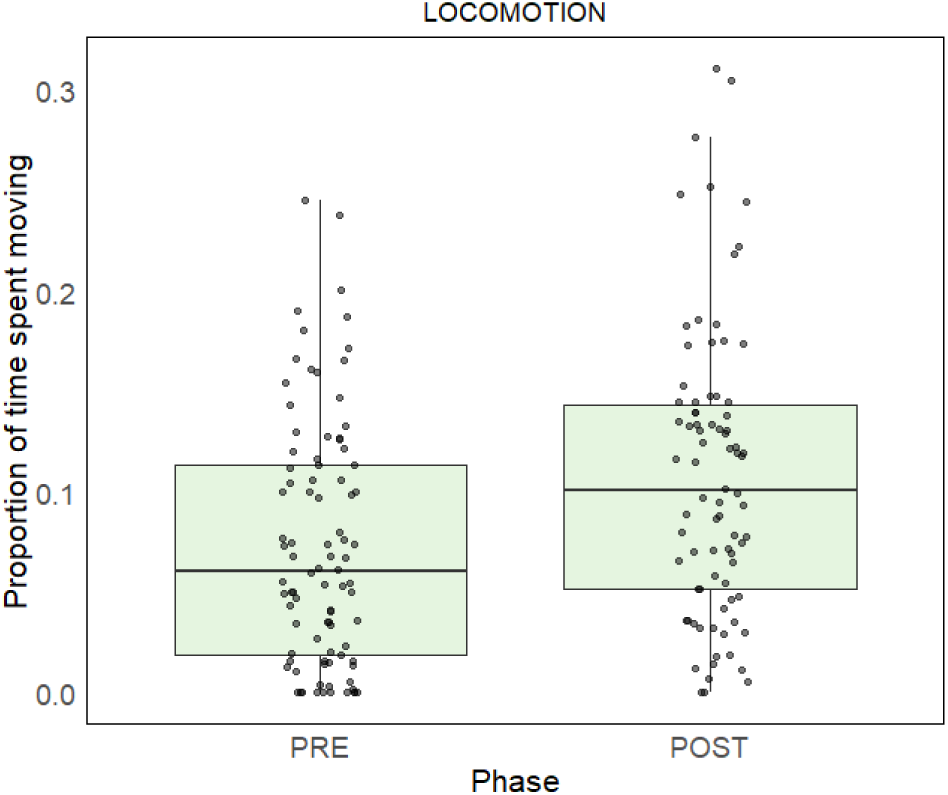
Proportion of time spent moving in pre-treatment and post-treatment phases.

In Model 4 (object manipulation) the full vs. null model comparison was not significant (χ2= 5.433, df=3, p=0.143), therefore post□hoc comparisons were not performed. Similarly, in Model 5 (abnormal behaviours) the full vs. null model comparison was not significant (χ2= 1.749, df=3, p= 0.63), therefore post□hoc comparisons were not performed. In Model 6 (cortisol levels) the full vs. null model comparison was significant (χ2= 16.908, df=3, p<0.001) and post□hoc comparisons indicated that cortisol values were lower in the post-treatment phase compared to the pre-treatment phase (β = -1.396, SE= 0.361, p=0.001, Figure 3). Full model outputs are reported in supplementary materials S-2.

**Figure 3.**
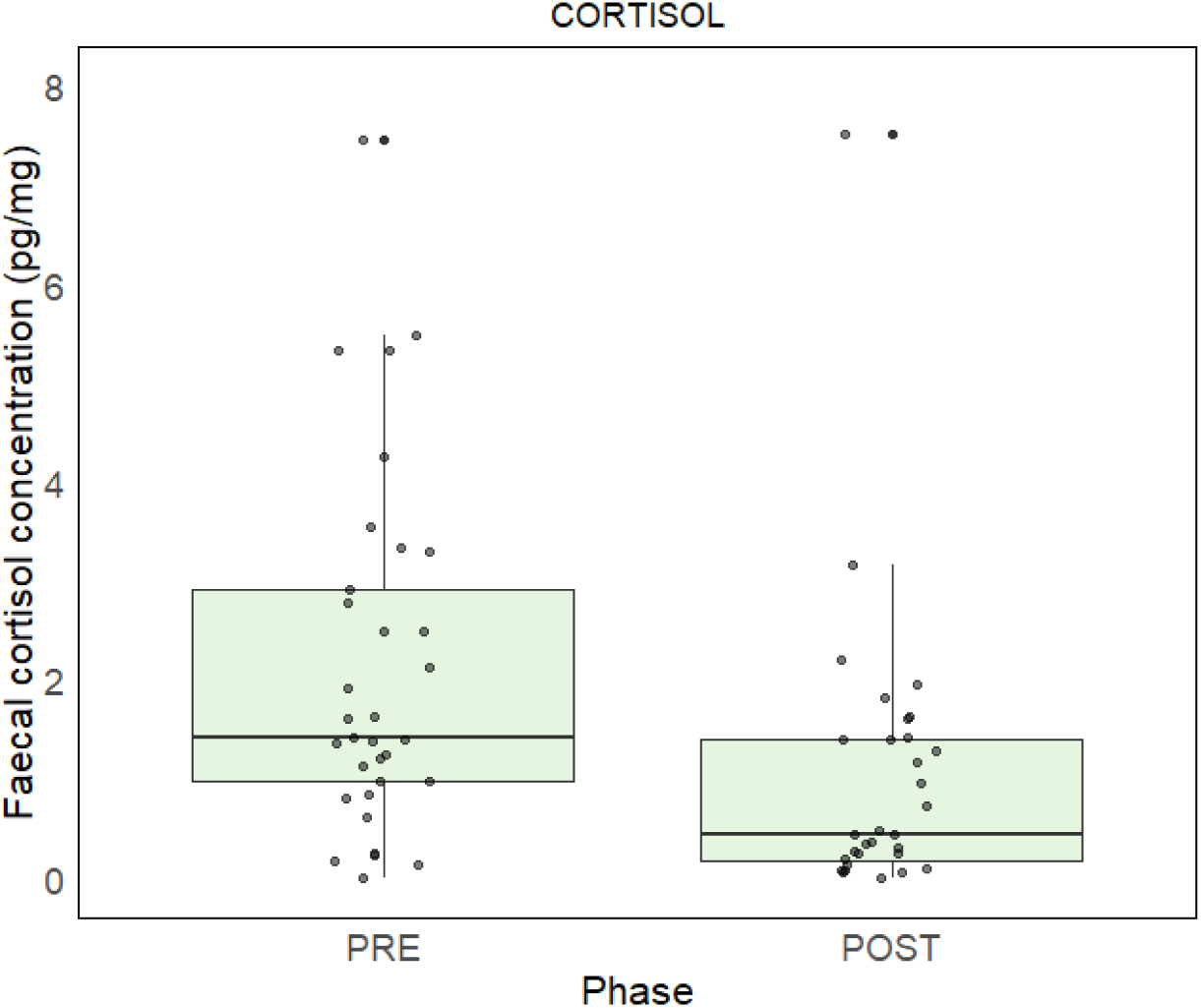
Faecal cortisol concentration in pre-treatment and post-treatment phases.

## 4. DISCUSSION

We tested whether environmental enrichment could have beneficial effects on the welfare of three aged macaques (*Macaca mulatta*) previously housed under laboratory conditions. We found partial evidence to support Prediction 1: there is evidence of an increase in locomotion (prediction 1a), but not objects manipulation (prediction 1b), in the post-treatment phase compared to the pre-treatment phase. Similarly, we found partial evidence to support Prediction 2: there is evidence of a decrease in scratches and body-shaking in the post-treatment phase compared to the pre-treatment phase, but we observed no significant difference in rates of abnormal behaviours between pre-treatment and post-treatment phase. Finally, in accordance with Prediction 3, we found a decrease in faecal cortisol concentration in the post-treatment phase compared to the pre-treatment phase.

The present study was designed to investigate the behavioural patterns of three aged rhesus macaques (≥ 20 years), following their relocation from a laboratory setting to the Animanatura Wild Sanctuary in Italy, and to investigate the potential benefits of environmental enrichment on the welfare of the subjects. Rhesus macaques are frequently utilised as non-human primate models in various domains of biomedical research, thereby underscoring the imperative to ensure their welfare in captive environments (Padrell et al., 2021). We found a positive impact of enrichment on activity budgets of our subjects: locomotion increased in the post-treatment phase compared to the pre-treatment phase. Similarly, we observed a decrease in resting behaviour in both the enrichment and post-treatment phase compared to the pre-treatment phase. Our findings align with previous research demonstrating the benefits of environmental enrichment for captive primates. For example, studies on long-tailed macaques and ring-tailed lemurs have shown that enrichment can promote positive changes in activity budgets (Albanese et al., 2021; Lameris et al., 2021). Contrary to our prediction, we did not observe a significant difference in objects manipulation between the pre- and post-treatment phase. Our result contrast with previous studies on the subject, that showed a positive effect of novel food enrichment on object manipulation behaviour (Clark, 2017; Padrell et al. 2022). For example, Clark and Smith (2013) observed that chimpanzees exhibited more problem-solving behaviours and spent more time engaged in social play when a puzzle feeder was present. One possible explanation for our result could be drawn when considering the age of our subjects: elder individuals tend to be less curious and active than younger individuals (Watowich et al., 2020; Whatley et al., 2025), and they spend significantly less time engaging in object manipulation than babies and juveniles (Vauclair et al., 1994). Nonetheless, one concurrent cause that may have affected our result could reside in the housing setting: social facilitation between individuals plays a role in the reactions of primates to objects, and socially housed rhesus macaques were found to extensively manipulate objects even at an old age (Novak et al., 1993).

We found evidence of a positive effect of enrichment in reducing scratching and body shaking behaviour. This aligns with previous research demonstrating similar reductions in stress-related behaviours with the implementation of enrichment strategies (Lutz et al., 2022, Lutz and Baker, 2023). Our results suggests that enrichment can effectively mitigate stress and improve the overall well-being of aged captive primates, particularly those previously subjected to less stimulating environments such as laboratory cages. Specifically, studies on single housed rhesus macaques have shown that enrichment, including feeding enrichment, leads to a reduction in abnormal behaviours and an increase in species-typical behaviours such as play (Schapiro & Bloomsmith, 1995). Similarly, research on Javan gibbons has demonstrated that enrichment, decreased abnormal behaviours and encouraged more natural foraging activities (Gronqvist et al., 2013). Additionally, a recent study on ring-tailed lemurs showed a positive effect of enrichment in reducing rates of stress-related behaviours such as scratching (Caselli et al., 2022). Contrary to our prediction, we did not find a significant reduction in the proportion of abnormal behaviours in the post-treatment phase compared to the pre-treatment phase. These results could be attributed by the low number of instances of abnormal behaviours (n=4) observed in our subjects: only 3 instances of abnormal behaviour were observed in the pre-treatment phase, whereas we observed only one instance of abnormal behaviour in the post-treatment phase. Although our descriptive results support the positive impact of environmental enrichment on reducing abnormal behaviours in captive non-human primates, our sample size and the low number of observations are not sufficient to support our prediction from a statistical standpoint.

Finally, we observed a reduction of faecal cortisol concentration in the post-treatment phase compared to the pre-treatment phase. Our finding is consistent with several studies that previously indicated that successful enrichment programmes can lead to a decrease in faecal glucocorticoid metabolite concentrations in captive primates, suggesting a reduction in physiological stress (Boinski et al., 1999; Pizzutto et al., 2008; Vaglio et al., 2021, Richardson, 2024). Furthermore, the observed decrease in faecal cortisol levels in the post-treatment phase further supports the stress-reducing effects of enrichment (Wooddell et al., 2019). Nonetheless this cortisol-related results should be interpreted with caution, as various factors, including seasonality, can influence cortisol levels. However, the observed decrease in cortisol, combined with positive effects on behaviours, strongly supports the positive impact of enrichment in reducing stress and enhancing animal welfare (Keay et al., 2006). While some initial stress responses to novelty might occur (Albanese et al., 2021), the long-term effects of enrichment generally point to lower FCM levels, particularly when the enrichment promotes species-typical behaviours (Gronqvist et al., 2013).

Rehabilitation programmes should understand and consider the species-specific physiological, psychological, and behavioural needs of the animals to ensure their welfare while in rescue centres. In Italy, two pilot rehabilitation programmes for colonies of long-tailed macaques (*Macaca fascicularis*) from laboratories are currently active, aiming to create a model for the recovery of primates from labs (Albanese et al., 2021). In conclusion, the study underscores the importance of environmental enrichment as a means of improving the welfare and promoting species-appropriate behaviour in captive primates (Costa et al., 2018; Richardson, 2024), and these observations align with the notion that environmental enrichment can promote opportunities to develop species-specific behaviours and mitigate undesirable ones (Richardson, 2024). Additionally, these findings provide compelling evidence that the implementation of an enrichment program can significantly enhance the welfare of aged non-human primates and may be considered a valuable tool in the rehabilitation of non-human primates with a history of being housed in laboratory settings for their entire life. While the current study provides encouraging evidence for the welfare-enhancing effects of environmental enrichment, the small sample size limits the generalizability of the findings. Future research with larger sample sizes and a more comprehensive assessment of behavioural and physiological indicators of welfare would further elucidate the potential benefits of enrichment for aged primates recovered from laboratory conditions.

## 5. ANIMAL WELFARE IMPLICATIONS

The findings of this study highlight the positive welfare implications of providing environmental enrichment to aged rhesus macaques with a history of laboratory housing. Overall, enrichment contributed to improved behavioural and physiological indicators of well-being, including increases in activity levels, reductions in stress-related behaviours such as scratching and body-shaking, and lower faecal cortisol concentrations. Although changes in object manipulation and abnormal behaviours were less pronounced—partly due to low baseline occurrence—the general pattern supports the view that enrichment can help mitigate stress and promote species-typical behaviours, even in older individuals. These results reinforce the importance of tailored enrichment programmes in rehabilitation settings, particularly for primates transitioning from restrictive laboratory environments, and underline the value of enrichment as a practical tool for enhancing welfare in captive, aged non-human primates.

## AUTHOR CONTRIBUTION

VA, MK and RB performed data collection and data decoding. PA and GA performed cortisol analyses. FD and MP performed data analysis. Hormonal analyses were performed by GM. Manuscript preparation was led by FD and ML.

## ACKNOWLEDGMENTS

We thank LAV ETS and Animanatura Wild Sanctuary for allowing us to conduct this research project.

## DATA AVAILABILITY

The data that support the findings of this study are available on request.

