## Supplementary Materials for "Improving Welfare Through Enrichment: A Case Study in Aged Ex-Laboratory Rhesus Macaques"

**S-1 – STUDY SUBJECTS**

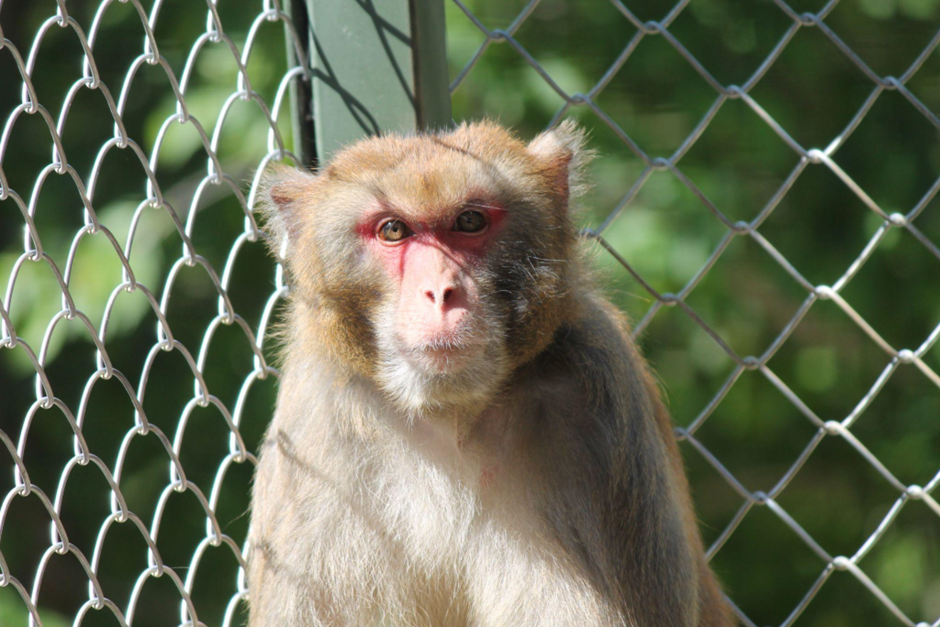

| **NAME** | Bob |
| --- | --- |
| **SEX** | M |
| DATE OF BIRTH (PRESUMED) | 02/01/2001 |

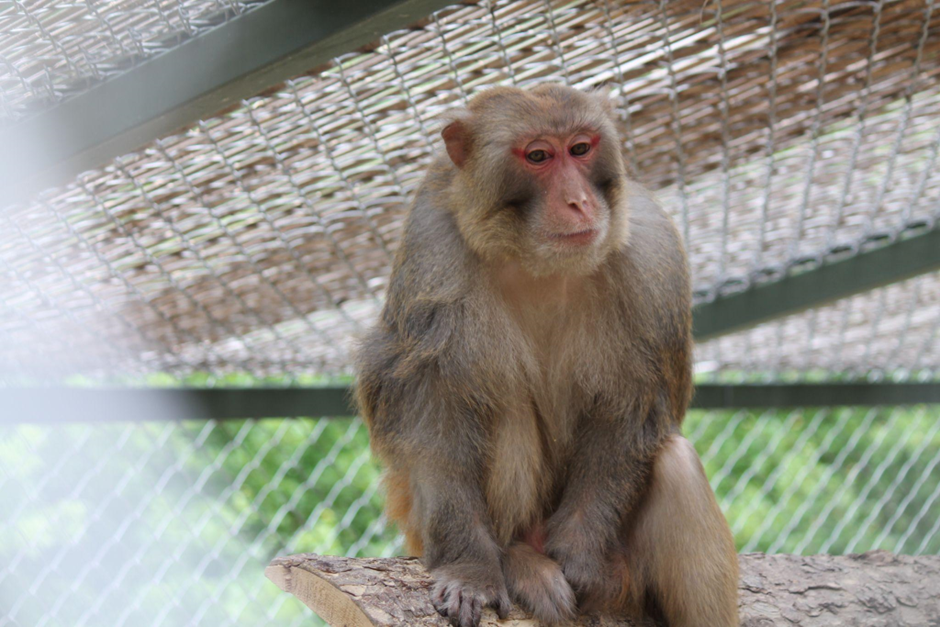

| **NAME** | Lucio |
| --- | --- |
| **SEX** | M |
| **DATE OF BIRTH (PRESUMED)** | 04/01/2001 |

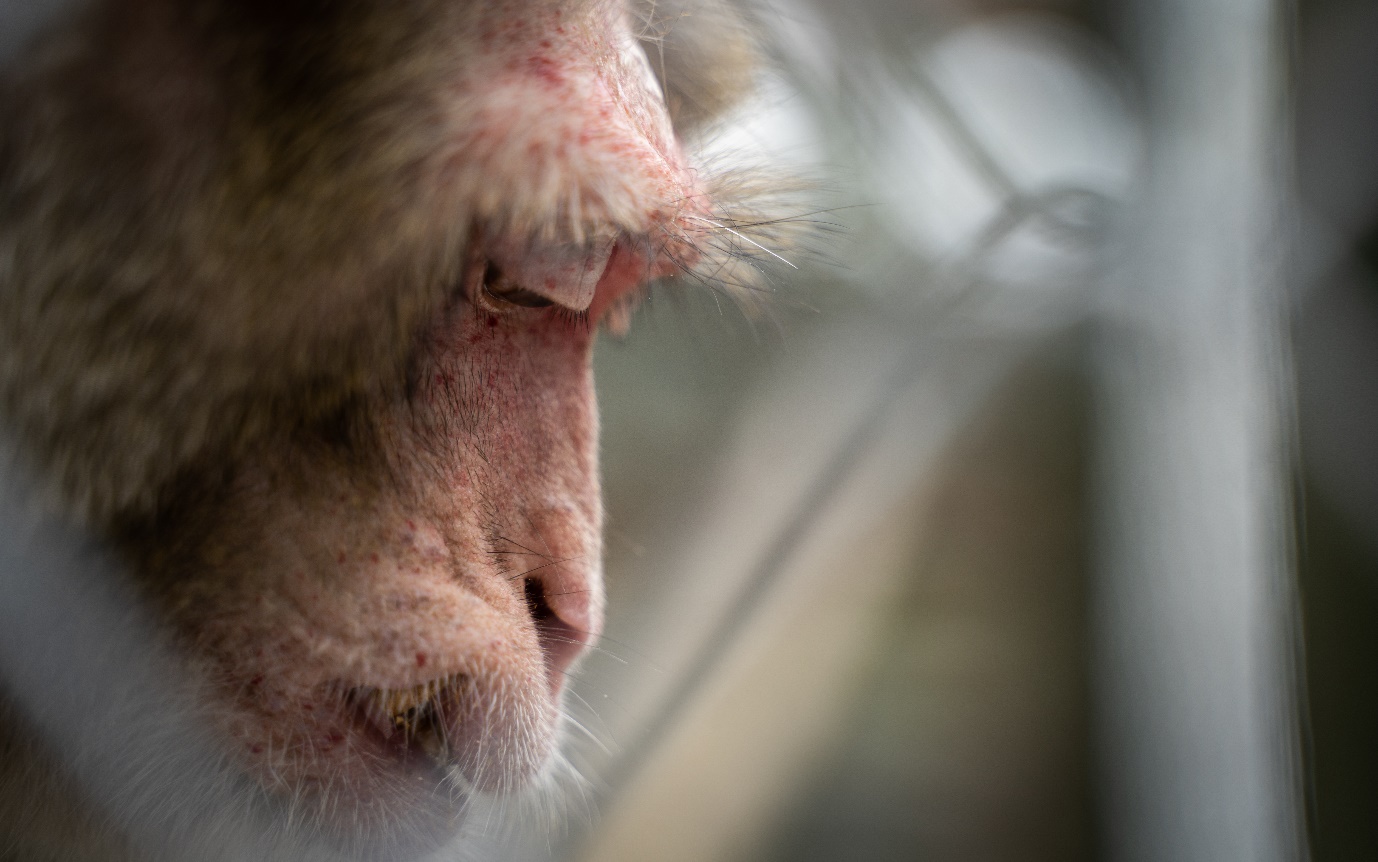

| **NAME** | Charlie |
| --- | --- |
| **SEX** | M |
| **DATE OF BIRTH (PRESUMED)** | 01/01/1993 |

**S-2 – ENVIRONMENTAL ENRICHMENTS AND ENCLOSURES**

**Enrichment activity/device**

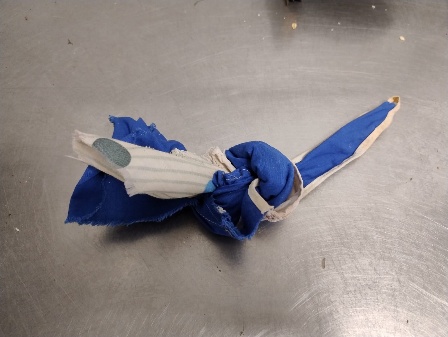

Cloth with dry fruit (20 gr)

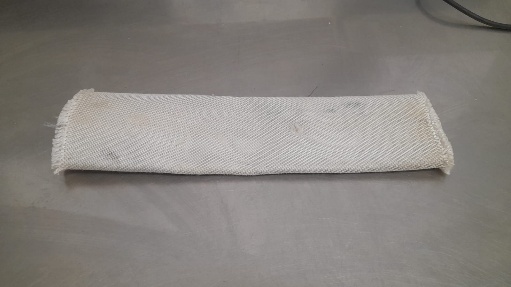

Firehose with boiled rice (30 cm length, 50 gr)

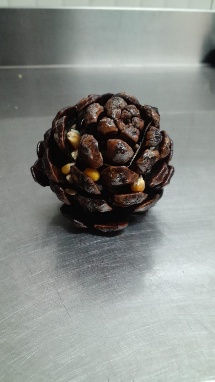

Pinecone with seeds

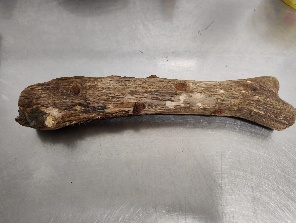

Stick with holes filled with dry fruits

Kong with jam

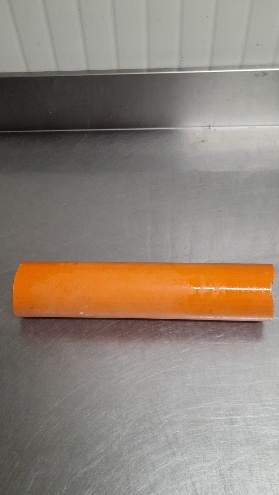

Pvc tube with honey and cornflakes (20 gr)

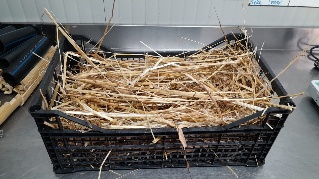

Plastic box with hay and seeds (50 gr)

Tab. S1 – Types of food enrichments used in the study.

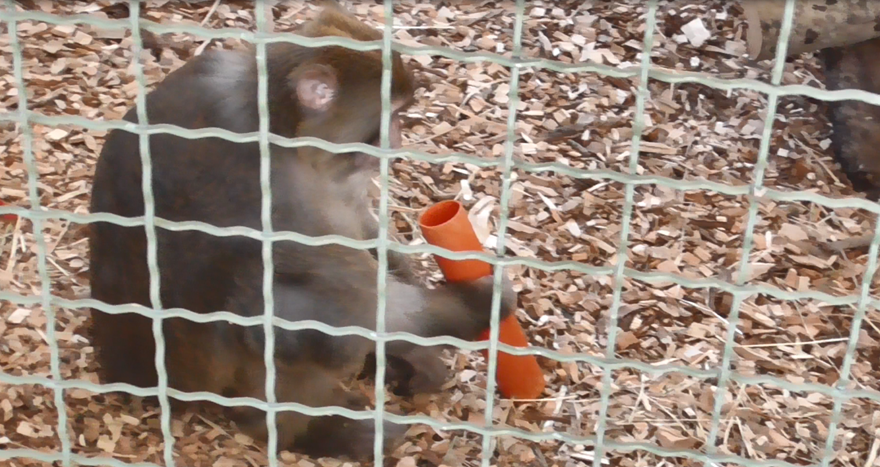

Fig. S-2. Example of enrichment session using Pvc tube with honey and cornflakes.

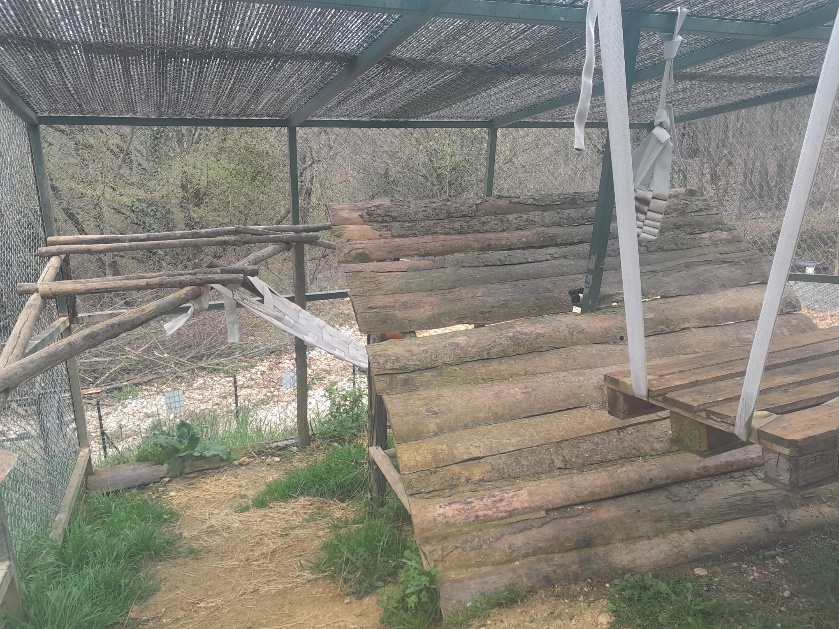

Fig. S-4. Interior view of the external enclosure.

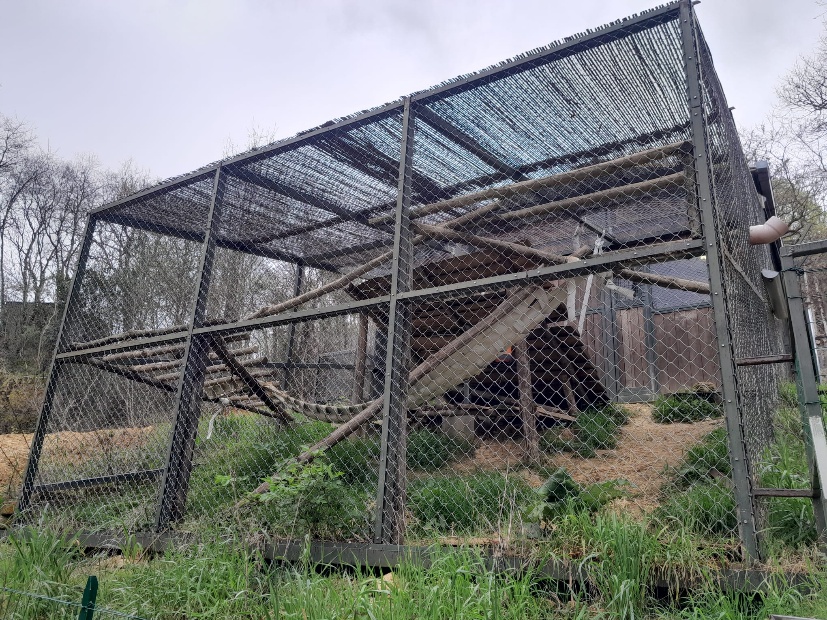

Fig. S-4. Exterior view of the external enclosure.

**S-3 – FULL MODEL OUTPUTS**

**MODEL 1**

**Table S-3.1** Results of the model 1, investigating body-shake behaviour (significant results are highlighted in bold)

| **Predictor** | **β** | **SE** | **p** |
| --- | --- | --- | --- |
| intercept | -2.748 | 0.896 | 0.002 |
| Phase treatment 2 | 0.013 | 0.397 | 0.973 |
| Phase post-treatment | -0.272 | 0.421 | 0.517 |
| Phase pre-treatment | 0.481 | 0.330 | 0.145 |
| Tourist presence | 0.63805 | 0.29677 | **0.031** |
| Max. daily temperature | -0.05862 | 0.02656 | **0.027** |
| Time of observation | -0.22475 | 0.20174 | 0.265 |

Full vs. null model: χ2=8.5089, df=3, p=0.037

**MODEL 2**

**Table S-3.2** Results of the model 2, investigating scratching behaviour (significant results are highlighted in bold)

| **Predictor** | **β** | **SE** | **p** |
| --- | --- | --- | --- |
| intercept | -3.275 | 0.642 | >0.001 |
| Phase treatment 2 | 0.449 | 0.230 | 0.051 |
| Phase post-treatment | 0.497 | 0.249 | **0.046** |
| Phase pre-treatment | 0.990 | 0.200 | **>0.001** |
| Tourist presence | 0.098 | 0.221 | 0.657 |
| Max. daily temperature | 0.003 | 0.019 | 0.840 |
| Time of observation | -0.012 | 0.115 | 0.915 |

Full vs. null model: χ2=32.479, df=3, p<0.001

**MODEL 3**

**Table S-3.3** Results of the model 3, investigating locomotion behaviour (significant results are highlighted in bold).

| **Predictor** | **β** | **SE** | **p** |
| --- | --- | --- | --- |
| intercept | -2.312 | 0.466 | >0.001 |
| Phase treatment 2 | 0.346 | 0.127 | **0.006** |
| Phase post-treatment | 0.284 | 0.168 | 0.091 |
| Phase pre-treatment | -0.208 | 0.136 | 0.125 |
| Tourist presence | -0.019 | 0.197 | 0.923 |
| Max. daily temperature | -0.004 | 0.013 | 0.740 |
| Time of observation | 0.206 | 0.088 | **0.019** |

Full vs. null model: χ2= 21.55, df=3, p<0.001

**MODEL 4**

**Table S-3.4** Results of the model 4, investigating object manipulation behaviour (significant results are highlighted in bold).

| **Predictor** | **β** | **SE** | **p** |
| --- | --- | --- | --- |
| intercept | -3.949 | 0.528 | >0.001 |
| Phase treatment 2 | 0.068 | 0.145 | 0.636 |
| Phase post-treatment | -0.251 | 0.191 | 0.190 |
| Phase pre-treatment | -0.209 | 0.146 | 0.153 |
| Tourist presence | -0.018 | 0.237 | 0.937 |
| Max. daily temperature | -0.004 | 0.015 | 0.780 |
| Time of observation | -0.001 | 0.101 | 0.986 |

Full vs. null model: χ2= 5.433, df=3, p=0.143

**MODEL 5**

**Table S-3.5** Results of the model 5, investigating abnormal behaviours (significant results are highlighted in bold).

| **Predictor** | **β** | **SE** | **p** |
| --- | --- | --- | --- |
| intercept | -5.707 | 0.348 | >0.001 |
| Phase treatment 2 | -0.045 | 0.095 | 0.635 |
| Phase post-treatment | -0.140 | 0.127 | 0.269 |
| Phase pre-treatment | -0.001 | 0.094 | 0.990 |
| Tourist presence | -0.017 | 0.010 | 0.092 |
| Max. daily temperature | 0.337 | 0.134 | **0.011** |
| Time of observation | -0.026 | 0.064 | 0.685 |

Full vs. null model: χ2= 1.749, df=3, p= 0.626

**MODEL 6**

**Table S-3.6** Results of the model 6, investigating faecal cortisol levels (significant results are highlighted in bold).

| **Predictor** | **β** | **SE** | **p** |
| --- | --- | --- | --- |
| intercept | 1.538 | 1.275621 | 0.227874 |
| Phase treatment 2 | -0.576 | 0.301 | 0.055 |
| Phase post-treatment | -1.531 | 0.421 | **0.0002** |
| Phase pre-treatment | -0.196 | 0.334 | 0.556 |
| Tourist presence | 1.022 | 0.443 | **0.021** |
| Max. daily temperature | 0.004 | 0.038 | 0.912 |
| Time of observation | -0.561 | 0.220 | **0.010** |

Full vs. null model: χ2= 16.908, df=3, p<0.001

**S-4 – POST-HOC COMPARISONS**

Post-hoc comparisons are shown only for those models that showed a significant likelihood ratio test (i.e., a significant full vs null model comparison). In all models, estimates are in logit‐scale.

**Table S-4.1 -** Results of the post-hoc comparison of model 1, investigating body-shake behaviour (significant results are highlighted in bold).

| **Contrast** | **estimate** | **SE** | **p** |
| --- | --- | --- | --- |
| Treatment phase 1 - Treatment phase 2 | -0.013 | 0.398 | 1.000 |
| Treatment phase 1 – Post treatment | -0.420 | 0.421 | 0.750 |
| Treatment 1 - Pre-treatment | -1.174 | 0.330 | **0.002** |
| Treatment phase 2 - Post treatment | -0.407 | 0.375 | 0.699 |
| Treatment phase 2 - Pre-treatment | -1.161 | 0.307 | **0.001** |
| Post treatment - Pre-treatment | -0.754 | 0.295 | **0.052** |

**Table S-4.2 -** Results of the post-hoc comparison of model 2, investigating scratching behaviour (significant results are highlighted in bold).

| **Contrast** | **estimate** | **SE** | **p** |
| --- | --- | --- | --- |
| Treatment phase 1 - Treatment phase 2 | -0.450 | 0.231 | 0.207 |
| Treatment phase 1 – Post treatment | -1.191 | 0.250 | **<0.0001** |
| Treatment 1 - Pre-treatment | -1.683 | 0.200 | **<0.0001** |
| Treatment phase 2 - Post treatment | -0.741 | 0.205 | **0.002** |
| Treatment phase 2 - Pre-treatment | -1.234 | 0.169 | **<0.0001** |
| Post treatment - Pre-treatment | -0.493 | 0.171 | **0.020** |

**Table S-4.3 -** Results of the post-hoc comparison of model 3, investigating locomotion behaviour (significant results are highlighted in bold).

| **Contrast** | **estimate** | **SE** | **p** |
| --- | --- | --- | --- |
| Treatment phase 1 - Treatment phase 2 | -0.346 | 0.127 | **0.032** |
| Treatment phase 1 – Post treatment | -0.284 | 0.169 | 0.330 |
| Treatment 1 - Pre-treatment | 0.208 | 0.136 | 0.418 |
| Treatment phase 2 - Post treatment | 0.061 | 0.142 | 0.972 |
| Treatment phase 2 - Pre-treatment | 0.555 | 0.128 | **0.0001** |
| Post treatment - Pre-treatment | 0.493 | 0.150 | **0.005** |

**Table S-4.4 -** Results of the post-hoc comparison of model 6, investigating faecal cortisol levels (significant results are highlighted in bold).

| **Contrast** | **estimate** | **SE** | **p** |
| --- | --- | --- | --- |
| Treatment phase 1 - Treatment phase 2 | 0.563 | 0.312 | 0.272 |
| Treatment phase 1 – Post treatment | 1.362 | 0.444 | **0.011** |
| Treatment 1 - Pre-treatment | -0.034 | 0.337 | 0.999 |
| Treatment phase 2 - Post treatment | 0.799 | 0.799 | 0.199 |
| Treatment phase 2 - Pre-treatment | -0.597 | 0.303 | 0.198 |
| Post treatment - Pre-treatment | -1.396 | 0.361 | **0.001** |
